# FTO separation-of-function mutations alter m^6^A versus m^6^A_m_ demethylation selectivity on RNA

**DOI:** 10.64898/2026.05.19.726201

**Authors:** Kasun Dilshan Abeyrathne Eluwawalage, Brittany Shimanski, Marcin Warminski, Suma Katta, Rebecca Payne, Yanbao Yu, Joanna Kowalska, Jacek Jemielity, Jeffrey S. Mugridge

## Abstract

The RNA demethylase FTO erases N6-methyladenosine (m^6^A) and cap-associated N6,2′-O-dimethyladenosine (m^6^A_m_) modifications. However, the molecular basis of its substrate selectivity and the biological effects of m^6^A versus m^6^A_m_ demethylation in cells remain poorly understood. Here we report two engineered FTO separation-of-function mutants to selectively demethylate either m^6^A or m^6^A_m_ modifications on RNA. While investigating the propensity of FTO active site residues to undergo self-hydroxylation, we found that mutations of FTO residue L203 resulted in impaired m^6^A demethylation but retained wild-type levels of m^6^A_m_ demethylation, and that FTO L203A could function as a selective m^6^A_m_ demethylase. Conversely, building on our recent work that identified conserved aromatic residues on FTO involved in mRNA 5′ cap recognition, we found that the FTO H232A/W278A double mutant efficiently demethylates m^6^A modifications while exhibiting substantially impaired m^6^A_m_ demethylation, making it a selective m^6^A demethylase. Together, these complementary FTO variants represent the first set of engineered mutations that shift FTO demethylation selectivity between m^6^A and m^6^A_m_ substrates. These tools enable selective enzymatic removal of m^6^A or m^6^A_m_ modifications *in vitro* for sequencing applications, and may facilitate understanding of FTO-mediated m^6^A versus m^6^A_m_ demethylation in cellular and disease model systems.

## INTRODUCTION

Chemical modifications of RNA are important regulators of RNA function and stability and are closely linked to many human physiological processes and disease progression (Fang et al., 2021; He & He, 2021; R. Wu et al., 2016). N6-methyladenosine (m^6^A) is the most abundant and well characterized mRNA modification, with an average of 1 – 2 m^6^A modifications per 1000 nucleotides in eukaryotic mRNA (Hu et al., 2022; Song & Yi, 2020). These methyl modifications are enriched in the stop codon, 3’ untranslated regions (UTR), and in long internal exons (Dominissini et al., 2012; C. M. Wei & Moss, 1977). m^6^A participates in all steps of mRNA metabolism and function, including splicing, folding, export, translation, and degradation, thus impacting gene expression and diverse cellular processes (Lesbirel & Wilson, 2019; H. Liu et al., 2019; N. Liu et al., 2017; Q. Liu & Gregory, 2019; Mao et al., 2019; Yang et al., 2018; Zhu et al., 2023). The related modification, N6,2′-O-dimethyladenosine (m^6^A_m_), can be found at the first transcribed nucleotide as part of the 5′ cap structure in up to 30 % of eukaryotic mRNA transcripts and also at the 5′ end of some snRNAs (J. F. Liu et al., 2025; Mauer et al., 2019; C. M. Wei et al., 1975). Although the biological roles of m^6^A_m_ are less established compared to m^6^A, this 5′ end modification is thought to play roles in mRNA splicing, stability and translation (Wang et al. 2023; Boo and Kim 2020; Lee and Rio 2015; Goh et al. 2020; Mauer et al. 2016; Sun et al. 2021a).

m^6^A and m^6^A_m_ are thought to be reversible, dynamically regulated RNA modifications in cells. These N6 methyl groups are installed by ‘writer’ enzymes METTL3/14 (m^6^A) and PCIF1 (m^6^A_m_), and can be removed by ‘eraser’ enzymes FTO (m^6^A and m^6^A_m_) and AlkBH5 (m^6^A) (**Figure 1A**) (Wu et al. 2023; Kaur et al. 2022; Gao et al. 2024; Wei et al. 2018; Wang et al. 2016, 2022). The m^6^A/m^6^A_m_ writers are *S*-adenosyl methionine (SAM)-dependent methyltransferases with relatively well-defined substrate specificities – METTL3/14 methylates adenosines within DRACH consensus motifs and PCIF1 methylates cap-proximal adenosines (Boulias et al., 2019; Woodcock et al., 2019). However, substrate selectivity and its determinants for Fe(II)-dependent eraser enzymes are less defined, particularly for RNA demethylase FTO, which has been proposed to demethylate both m^6^A and m^6^A_m_ RNA modifications in different cellular contexts.(Mauer et al. 2017, 2019). FTO plays key roles in regulating mRNA splicing during adipogenesis, fat cell differentiation and metabolism, is critical during human development, and has been linked to various human cancers including glioblastoma, leukemia, breast, gastric and cervical cancer (Church et al., 2009; Dong et al., 2025; Kaklamani et al., 2011; Li et al., 2017; M. Zhang et al., 2015; Zhao et al., 2014; Zou et al., 2019). Hence, biochemical tools to selectively demethylate m^6^A_m_ versus m^6^A RNA modifications would be useful toward understanding FTO selectivity and the biological roles of FTO-mediated m^6^A_m_ versus m^6^A demethylation in the cell and in human disease progression.

**Figure 1.**
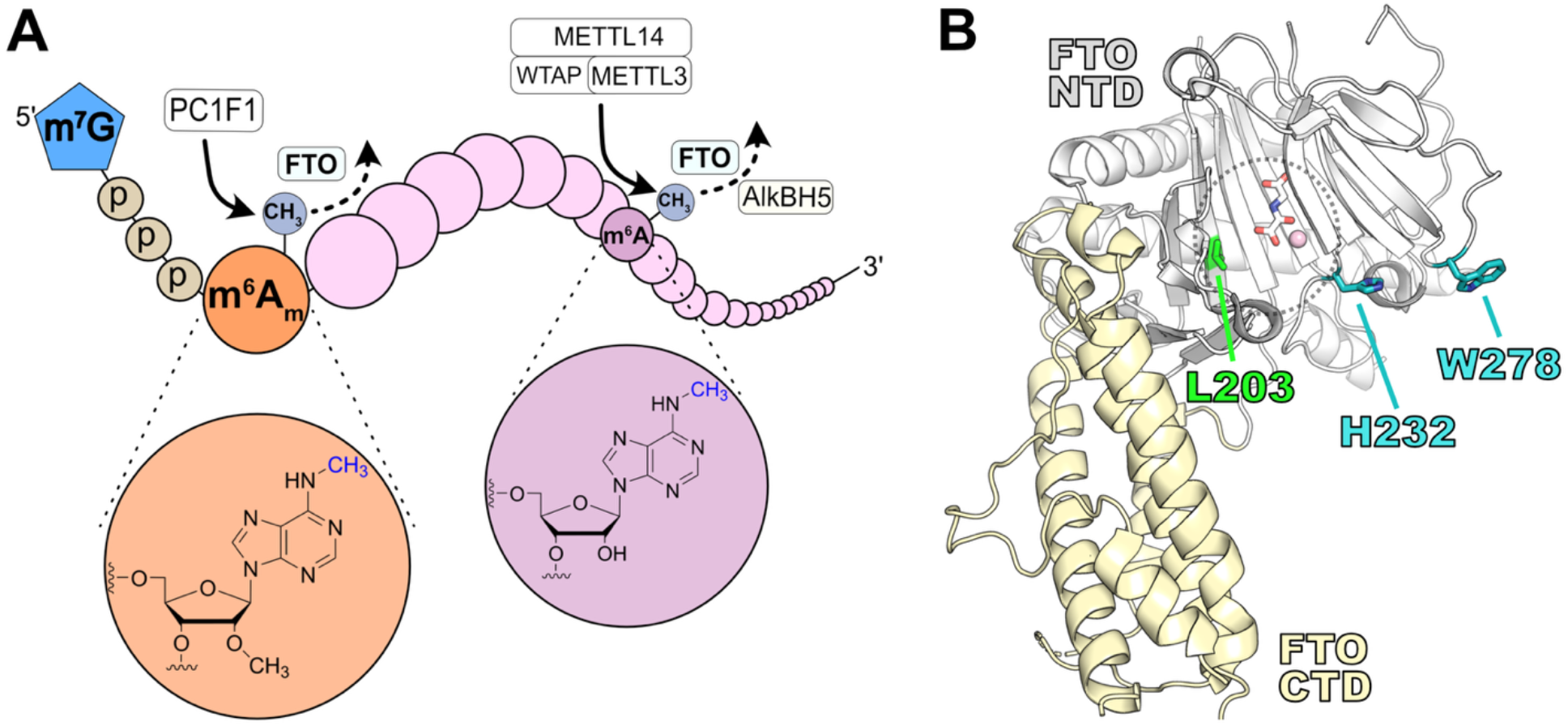
m^6^A and m^6^A_m_ RNA modifications and demethylase FTO. **(A)** Schematic of a canonical eukaryotic mRNA depicting internal m^6^A (pink) and cap-associated m^6^A_m_ (orange) modifications, along with their respective writer (METTL3/14, PCIF1) and eraser (FTO, ALKBH5) enzymes. **(B)** Structure of FTO (from PDB 5ZMD) showing the N-terminal catalytic domain (NTD, white) and C-terminal helical domain (CTD, yellow), with key sites of mutation in this study highlighted in green (L203) and teal (H232 and W278). The FTO active site is indicated with a dotted circle and includes bound 2-oxoglutarate analog (white sticks) and inactive metal Mn^2+^ (pink sphere).

In this study we report and characterize two FTO separation-of-function variants: FTO L203A, which selectively demethylates m^6^A_m_ but not m^6^A modifications on RNA, and FTO H232A/W278A, which selectively demethylates m^6^A over m^6^A_m_ modifications (**Figure 1B**). Together, these engineered FTO constructs represent the first set of mutations capable of shifting FTO’s demethylation selectivity between m^6^A and m^6^A_m_ substrates. These tools have dual utility: *in vitro* they enable selective demethylation of m^6^A or m^6^A_m_ for sequencing applications, and in cellular contexts they may allow researchers to differentiate the effects FTO-mediated m^6^A versus m^6^A_m_ demethylation in biology pathways and disease models.

## RESULTS

### FTO active site residue L203 is a site of self-hydroxylation and modulates FTO activity and selectivity

The human AlkBH family consists of nine homologous enzymes (AlkBH1-8 and FTO) that belong to the Fe(II)/2-oxoglutarate-dependent dioxygenase (Fe/2OG) superfamily, and carry out Fe(II)-mediated oxidation reactions on DNA or RNA to help maintain and regulate the epigenome and epitranscriptome(Das et al., 2025; Fu & Samson, 2012; Kaur et al., 2022; Kogaki et al., 2023; F. Liu et al., 2016; Van Den Born et al., 2011; L. Zhang et al., 2024; X. Zhang et al., 2019) An interesting feature observed for some of these enzymes is their propensity to undergo self-hydroxylation of active site residues. For example, Fe/2OG enzymes AlkB, TfdA, and TauD can all self-hydroxylate aromatic residues in their own active sites (Liu et al., 2001; Henshaw et al., 2004; Ryle et al., 2003). Most recently, DNA/RNA demethylase AlkBH3 was shown to self-hydroxylate an active site leucine residue, and mutagenesis of that site resulted in somewhat altered dm1A versus dm3C demethylation selectivity on ssDNA (Sundheim et al., 2006). Like AlkBH3, FTO has an active site Leu residue (L203) at the same position (**Figure 2A,B**), and we found using peptide mass spectrometry that FTO L203 can similarly undergo self-hydroxylation under a variety of reaction conditions (**Figure 2C**).

**Figure 2.**
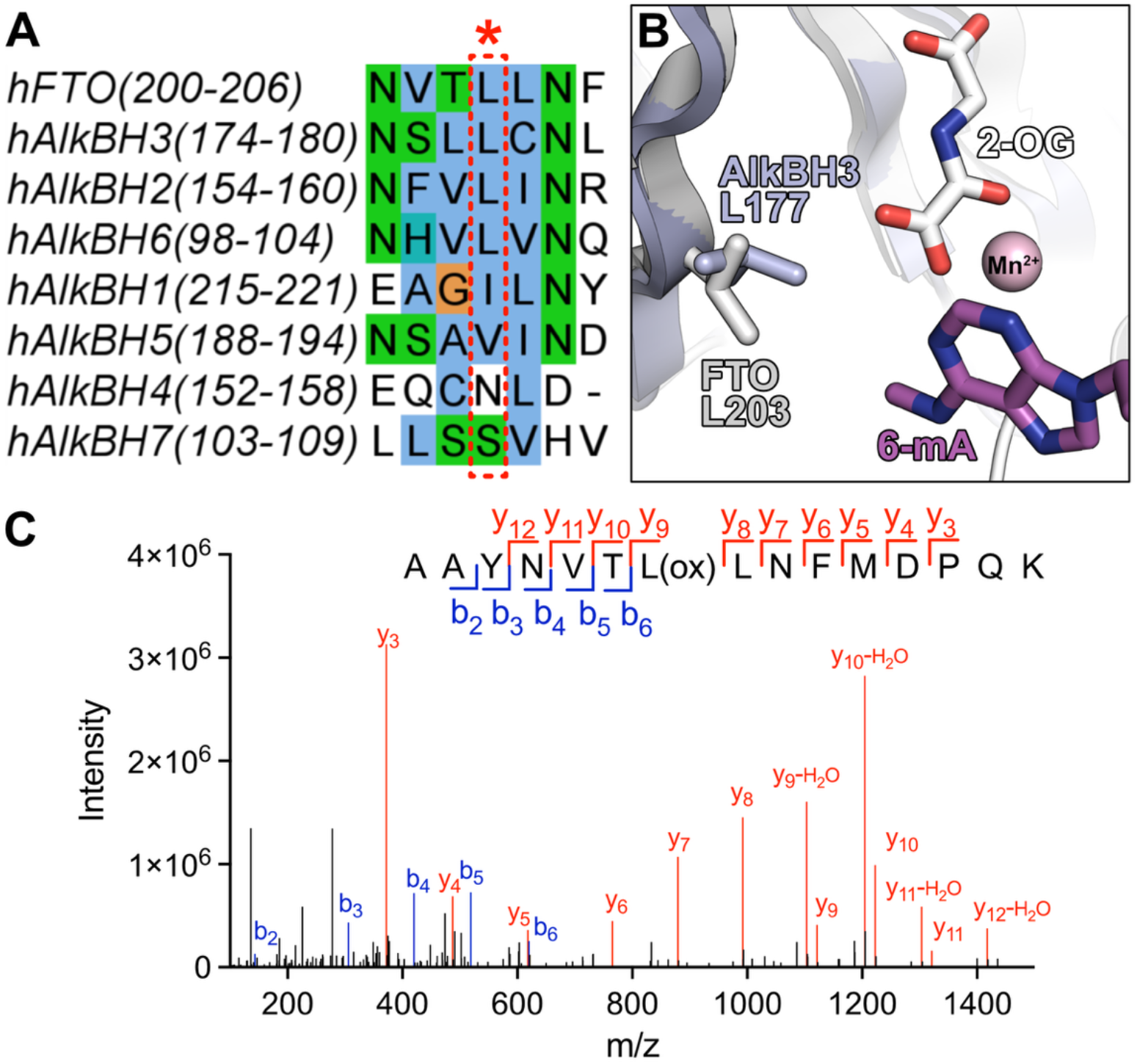
The FTO active site residue L203 undergoes self-hydroxylation. **(A)** Multiple sequence alignment of full length human AlkBH family members showing conservation of an active site leucine (red dotted box) in hFTO, hAlkBH3, hAlkBH2, and hAlkBH6. **(B)** Superposition of FTO (PDB 5ZMD, white) and AlkBH3 (PDB 9NCZ, slate) structures showing the positioning of conserved active site residues L203 (FTO) and L177 (AlkBH3) relative to the 6mA positioning in the FTO structure. **(C)** MS/MS spectrum showing L203 oxidation in the FTO tryptic peptide (197)-AAYNVTLLNFMDPQK-(211). The b and y ion series are highlighted in blue and red, respectively.

Toward understanding the function of FTO self-hydroxylation at position L203 and the possible roles of L203 in substrate selectivity, we next generated an L203X mutant panel to test how mutations at this position impact both m^6^A and m^6^A_m_ demethylation on model RNA substrates (**Figure 3**). FTO L203 mutations generally decreased both m^6^A and m^6^A_m_ demethylation activity, but in two cases, L203A and L203M, mutation decreased activity on internal m^6^A-containing substrates but not on 5′ m^6^A_m_-containing cap substrates; this was particularly true for the L203A mutation, which retained WT-level m^6^A_m_ demethylation activity but only ∼ 10% m^6^A demethylation activity. Although the function of leucine oxidation in the FTO and AlkBH3 active sites remains unclear and a topic for future study, these experiments suggested that the FTO L203A mutation could be a useful biochemical tool to selectively demethylate m^6^A_m_ modifications on RNA.

**Figure 3.**
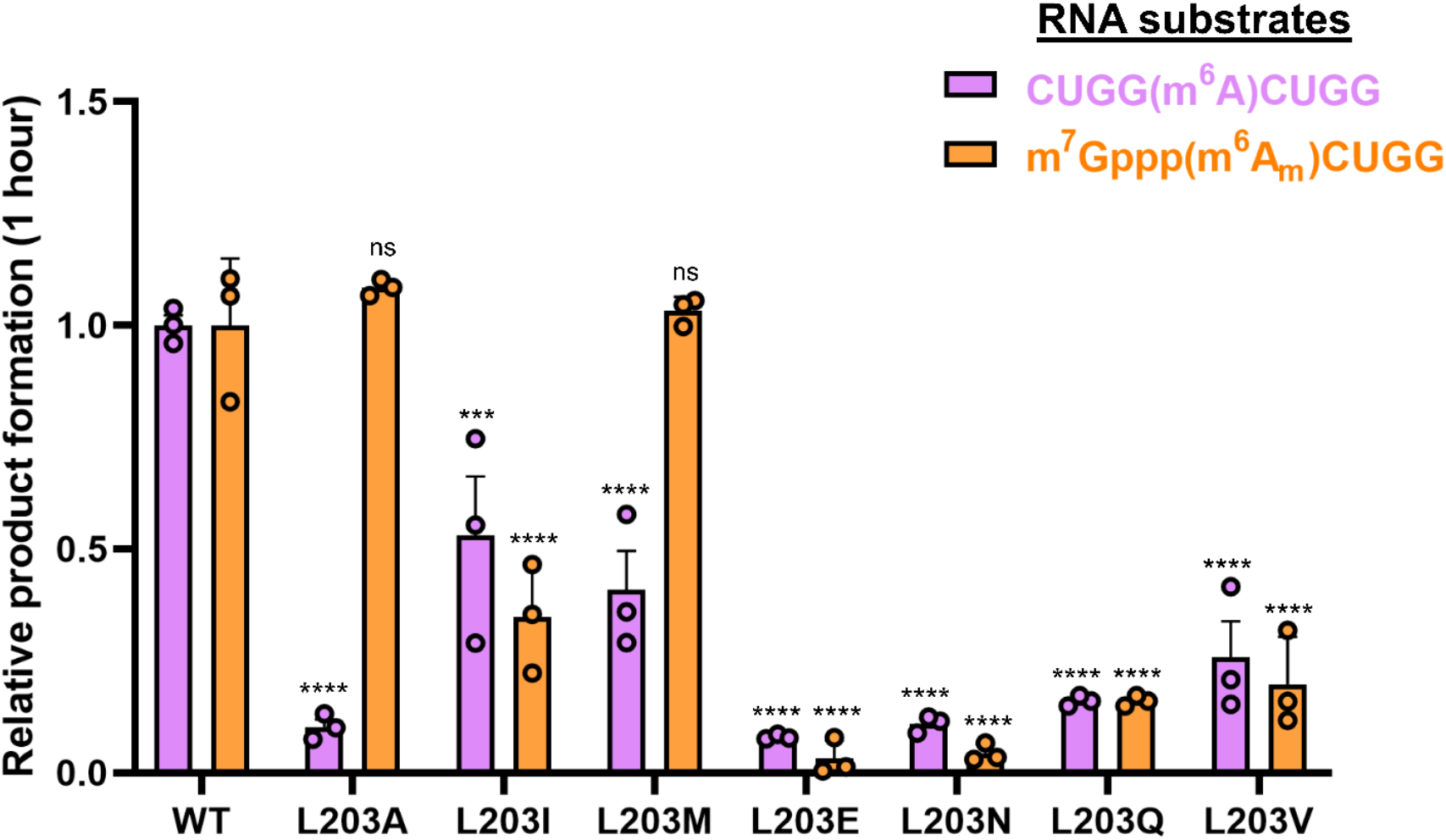
FTO L203A shows distinct demethylation behavior towards m^6^A or m^6^A_m_ model RNA substrates *in vitro*. Relative adenosine or 2′-O-methyladenosine product formation in demethylation assays with FTO L203X mutants with either CUGG(m^6^A)CUGG or m^7^Gppp(m^6^A_m_)CUGG model RNA substrates. Endpoint demethylation assays were conducted with 2 µM CUGG(m^6^A)CUGG or m^7^Gppp(m^6^A_m_)CUGG substrate and 1 µM FTO enzyme in triplicate. Product formation after 1 hour is normalized to WT activity. Data are shown as mean values ± SEM (n = 3) and statistical significance was calculated using a one-way ANOVA with Dunnett’s multiple comparisons test (****p<0.0001, ***p<0.001, ns p>0.05).

### FTO L203A selectively demethylates 5′ m^6^A_m_ modifications on RNA

To explore whether the FTO L203A mutant could be a useful biochemical tool for selective m^6^A_m_ demethylation on longer RNA substrates, we next carried out demethylation assays monitoring either formation of product A_m_ in reactions with a capped, m^6^A_m_-containing 40mer RNA substrate, or loss of m^6^A in reactions with an m^6^A-containing linear 25mer RNA or stem-loop 25mer RNA (**Figure 4**). Loss of m^6^A substrate is followed in these assays, rather than product adenosine formation, because the 25mer RNA substrate sequences contain multiple adenosines, making it difficult to monitor small changes in A concentration over the course of the reaction. Consistent with our results above, FTO L203A only showed significant demethylation activity towards the m^6^A_m_-containing substrate (**Figure 4A**) and did not show any appreciable activity with the m^6^A-containing linear or structured, stem-loop substrates (**Figure 4B,C**). These data suggest that the FTO L203A mutant can be used as a novel, engineered enzymatic tool to selectively remove m^6^A_m_ modifications, but not m^6^A modifications, on RNA.

**Figure 4.**
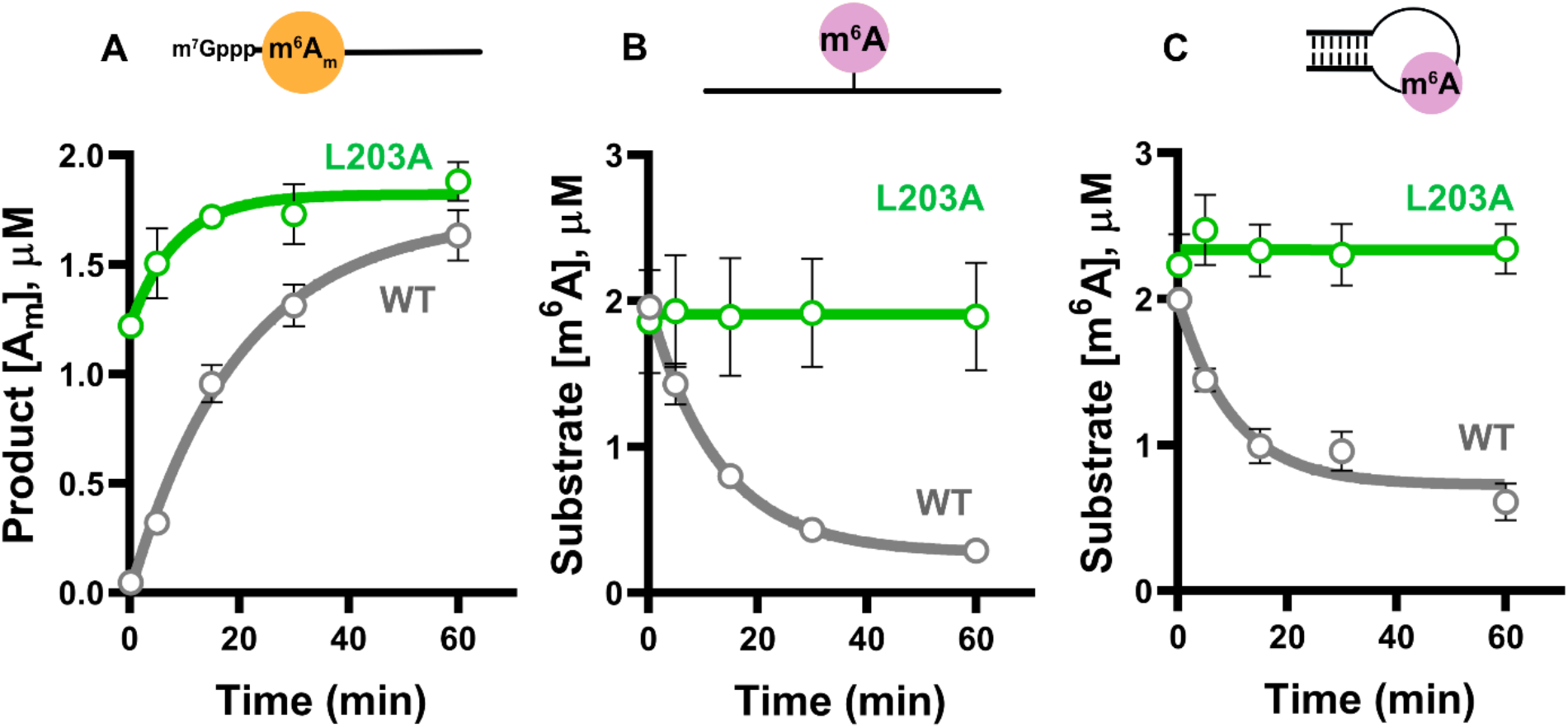
FTO L203A selectively demethylates m^6^A_m_-containing RNA. Time-course demethylation assays with wild-type FTO and FTO L203A using **(A)** a capped m^6^A_m_-containing 40mer RNA substrate, monitoring A_m_ product formation, **(B)** a linear m^6^A-containing 25mer RNA substrate, and **(C)** a structured stem-loop m^6^A-containing 25mer RNA substrate, monitoring loss of m^6^A substrate. Assays were conducted with 2 µM substrate and 1 µM enzyme in triplicate. Data points are shown as mean values ± SEM (n = 3).

### FTO H232A/W278A mutant selectively demethylates m^6^A modifications in RNA

We recently carried out hybrid molecular dynamics (MD) and biochemical studies to identify FTO residues that interact with the mRNA 5′ cap to promote m^6^A_m_ demethylation (Shimanski et al. 2025). This work identified two conserved, aromatic residues on the surface of FTO (H232 and W278; **Figure 1B**) that make stable, extended contacts by MD with the m^7^G group of the cap structure through cation-π and/or π-π interactions. Mutation of these aromatic FTO residues was found to impair m^6^A_m_ demethylation on model capped RNA oligos, but not on model m^6^A_m_-containing oligos lacking the m^7^G group, suggesting that these residues play a key role in specifically recognizing the mRNA 5′ cap structure. To further explore the utility of the FTO H232A/W278A double mutant as a tool for selective FTO-mediated m^6^A (but not m^6^A_m_) demethylation, we carried out demethylation reactions with FTO H232A/W278A and the capped, m^6^A_m_-containing 40mer RNA substrate and m^6^A-containing linear 25mer RNA or stem-loop 25mer RNA substrates (**Figure 5**), as above. These experiments showed that FTO H232A/W278A has a strong defect in m^6^A_m_ demethylation activity (**Figure 5A**), but nearly WT-level demethylation activity for m^6^A-containing substrates (**Figure 5B,C**). Although the m^6^A_m_ demethylation activity was not completely abolished by H232A/W278A double mutation, this FTO construct nevertheless has a dramatically shifted substrate preference (toward m^6^A) that could be used as a biochemical tool for selective FTO-mediated m^6^A demethylation.

**Figure 5.**
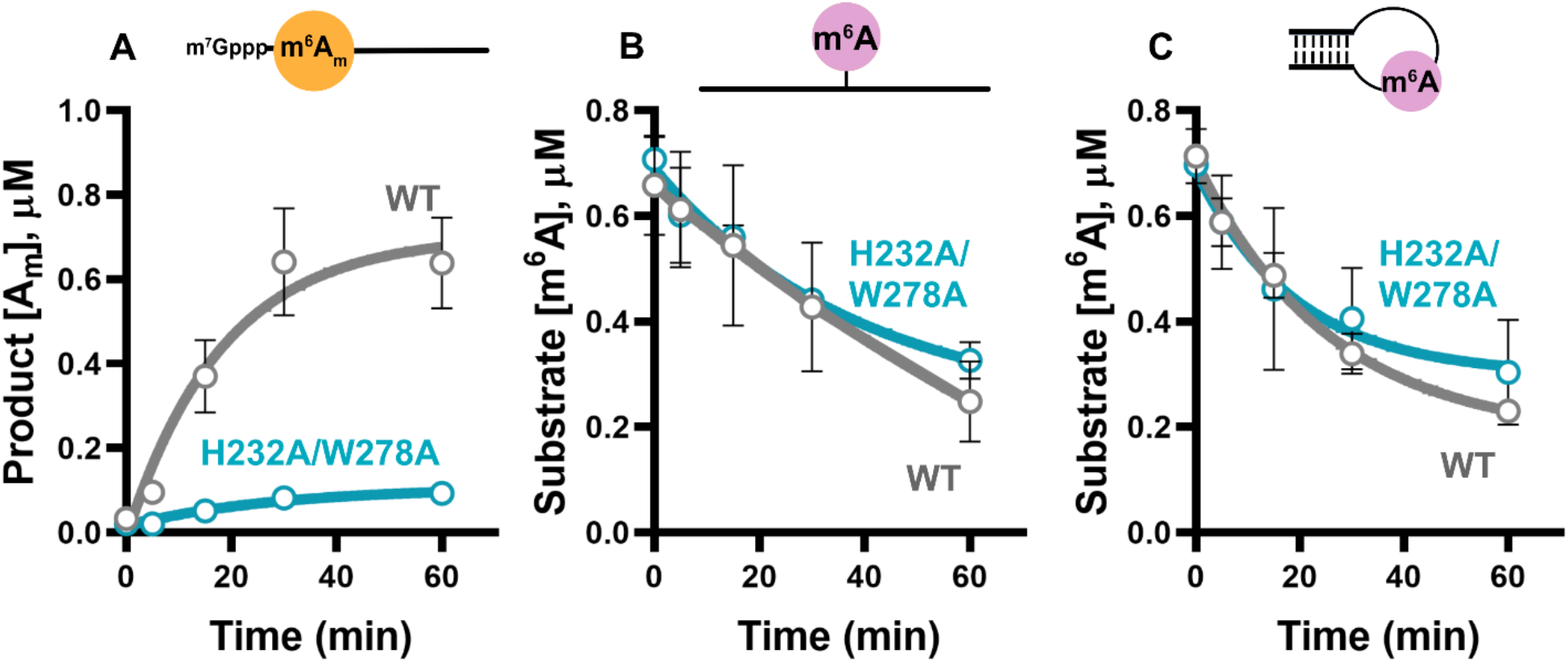
FTO H232A/W278A selectively demethylates m^6^A-containing RNA. Time-course demethylation assays with wild-type FTO and FTO H232A/W278A using **(A)** a capped m^6^A_m_-containing 40mer RNA substrate, monitoring A_m_ product formation, **(B)** a linear m^6^A-containing 25mer RNA substrate, and **(C)** a structured stem-loop m^6^A-containing 25mer RNA substrate, monitoring loss of m^6^A substrate. Assays were conducted with 0.25 µM enzyme and 0.75 µM substrate in triplicate. Data points are shown as mean values ± SEM (n = 3).

## DISCUSSION

FTO has been implicated in the demethylation of both internal m^6^A modifications on mRNA and cap-associated m^6^A_m_ modifications on mRNA and snRNA, but the molecular determinants governing its substrate selectivity remain incompletely understood. This has complicated efforts to assign specific biological functions or outcomes to FTO-mediated m^6^A versus m^6^A_m_ demethylation, particularly in cellular contexts where both modifications are present. In this study, we have characterized two engineered FTO variants with altered demethylation specificity: FTO L203A, which abolishes m^6^A demethylation activity while retaining activity on 5′ capped m^6^A_m_-containing substrates, and FTO H232A/W278A, which preferentially demethylates m^6^A relative to m^6^A_m_. Together, these separation-of-function variants provide a complementary set of biochemical tools for selective m^6^A versus m^6^A_m_ removal and for dissecting FTO-dependent RNA demethylation.

Our discovery that FTO L203A abolishes m^6^A demethylation activity without significantly affecting m^6^A_m_ activity was somewhat serendipitous – with characterization of FTO residue L203 originally motivated by the observation that other Fe/2OG family enzymes undergo self-oxidation of their active site residues. In particular, AlkBH3 active site residue L177 was previously shown to undergo self-hydroxylation and we found that the conserved Leu residue at this position in FTO (L203) also undergoes self-hydroxylation. Similar to AlkBH3, we found that mutation of L203 in FTO produced variable effects on the demethylation activity of different FTO substrates, with dramatic impairment of m^6^A demethylation, but not m^6^A_m_ demethylation, for the FTO L203A variant. In the FTO-6mA-DNA structure (PDB 5ZMD), L203 and 6mA methyl groups are located 4.3 Å apart, just outside the typical C-C van der Waals interaction distance; so while there does not appear to be a direct L203-m^6^A substrate contact, L203 may contribute to hydrophobic packing and the overall structuring of the FTO active site. From our data and the FTO-6mA-DNA structure, it remains mechanistically unclear why L203A mutation strongly impacts m^6^A but not m^6^A_m_ demethylation (after all, the N6-methyl groups would be expected to be similarly positioned in the active site in both cases), or what the likely effects and/or functions of L203 hydroxylation in FTO would be. Nevertheless, these investigations led to the discovery that the FTO L203A variant can selectively demethylate m^6^A_m_ but not m^6^A modifications on RNA.

In contrast, the H232A/W278A mutant provides an FTO variant with the opposite substrate bias: preferentially demethylating m^6^A over m^6^A_m_ modifications. These residues were identified through MD simulations in our recent report (Shimanski et al. 2025), and are proposed to specifically recognize the m^7^G group of mRNA 5′ cap. In this past study, we used only short, model RNA oligonucleotides, but show here that selective m^6^A demethylation with FTO H232A/W278A also persists on longer (25 – 40mer) RNA substrates. Although FTO H232A/W278A retains some minimal activity on m^6^A_m_-containing substrates, this mutation entirely reverses FTO’s intrinsic substrate preferences, making this engineered variant strongly biased toward m^6^A demethylation.

The two engineered FTO variants described here should be useful as biochemical tools for selective m^6^A versus m^6^A_m_ demethylation on RNA. Selective enzymatic removal of m^6^A_m_ by FTO L203A could aid workflows aimed at distinguishing cap-proximal m^6^A_m_ from internal m^6^A modifications, particularly in sequencing applications. A functionally similar approach that relies on using WT FTO in the absence of ascorbate was recently used to selectively remove m^6^A_m_ modifications for sequencing (Sun et al. 2021b), although our recent work shows that even in the absence of ascorbate, m^6^A modifications in structured RNAs can still undergo FTO-mediated demethylation (Calzini et al. 2025). Conversely, the FTO H232A/W278A variant, with its opposite substrate selectivity from WT FTO, may be useful for cell-based experiments to differentiate the biological effects of FTO-mediated m^6^A versus m^6^A_m_ demethylation in different cellular contexts or disease models. Together, these complementary, engineered separation-of-function variants show how FTO substrate selectivity is modulated by distinct active site and cap-binding residues, provide new biochemical tools for probing RNA modification biology, and may facilitate more precise interrogation of m^6^A/m^6^A_m_ epitranscriptomic regulation both *in vitro* and in cells.

## MATERIALS AND METHODS

### Protein expression and purification

*H. sapiens* FTO 32-505 DNA sequence was obtained as a single *E. coli* codon-optimized DNA sequence as a gBlock from Integrated DNA Technologies and cloned into a pET28a(+) kanamycin resistant expression vector with N-terminal 6x His tag. Single point mutants of FTO were introduced to the hFTO(32-505) construct using plasmid PCR site-directed mutagenesis. The vectors were transformed into *E. coli* BL21 (DE3) cells grown in LB media to OD ∼0.6 and protein expression was induced with IPTG for 18 h at 20 °C.

Cells were harvested at 5000 × g, lysed by sonication, and clarified at 16,000 × g in lysis buffer (25mM Tris-Base pH 8, 300mM NaCl, 5mM BME). The protein was purified by Ni-NTA affinity chromatography and buffer exchanged to (25mM Tris-HCl pH 7.5, 25mM NaCl). Solubility/affinity tags were cleaved by treatment with TEV protease overnight at 4 °C. FTO was further purified by size exclusion chromatography on a GE/Cytiva Superdex 200 16/60 gel filtration column in sizing buffer (10 mM HEPES pH 7, 150 mM NaCl, 1 mM DTT). The purified enzyme was concentrated to 30 mg/mL, flash frozen in liquid nitrogen, and stored at -70 ^°^C for future experiments.

The Edc1-Dcp1/Dcp2 decapping complex was purified according to the previously published procedure and stored at 4 mg/ml concentration (Paquette et al., 2018).

### m^6^A RNA substrate synthesis

CUGG(m^6^A)CUGG and m^6^A-25mer were purchased from Integrated DNA Technologies. Cap m^6^A_m_ RNA substrates (m^7^G_ppp_(m^6^A^m^)CUGG) were synthesized as previously described for similar RNA cap molecules. Oligonucleotide stemloop with the sequence 5′-5’-GGUUCCCGGUUGG(m^6^A)CUCCCGGGUUG-3′ containing the m^6^A (N6-methyladenosine) and 14th position was synthesized by using solid-phase phosphoramidites chemistry on a MerMade-4 Oligonucleotide synthesizer. The phosphoramidites were purchased from Glen research (PAC-rA, Ac-rC, rU, iPrPAC-rG) and 12 minute coupling time was used to couple m^6^A phosphoramidite and two such cycles were used. UltraMild Cap Mix A was used to prevent any exchange of Pac group with acetyl groups. The cleavage and deprotection step were carried out by treating the resin with aqueous methylamine at 65 ^°^C for 12 mins. 2’ O-TBDMS removal step was carried out with triethylamine and TEA.3HF. GlenPak cartridges were used to cleave off 5’ DMT group and the final concentration of oligonucleotide was determined by measuring UV absorbance at 260 nm.

### In vitro capped RNA generation

m^6^A_m_ capped RNA was transcribed according to previously published procedure with slight modifications (Warminski et al. 2024). A dsDNA template with class III promoter 6.5 containing only G at the +1-position was used. A large-scale IVT reaction (500 µL) was incubated overnight at 37 ^°^C (5 mM CTP, UTP, ATP nucleotides, 2 mM m^7^Gppp(m^6^A_m_)G, 0.05% triton-x, 5 mM DTT, 2 µM template, 0.05 mg T7 RNA polymerase, and 1 unit of TIPP) and cleaned up and concentrated with Zymo RNA clean up and concentrator kit. The final concentration of the capped oligoes was determined with a nanodrop by measuring the absorbance at 260 nm.

### *In vitro* m^6^A/m^6^A_m_ demethylation assays

2X FTO enzyme was incubated for 10 minutes in 1X reaction buffer (50 mM HEPES pH 7.0, 150 mM KCl, 100 µM (NH_4_)_2_Fe(SO_4_)2·6H_2_O) with cofactors (300 µM 2-oxoglutarate and 2 mM ascorbic acid); 2X RNA substrate was prepared in 1X reaction buffer. For demethylation assays with FTO H232A/W278A, final reaction mixtures contained 0.25 µM enzyme and 0.75 µM substrate, while all other demethylation assays were carried out with final concentrations of 1 µM enzyme and 2 µM substrate. Reactions were initiated by 1:1 addition of a 2X substrate mixture to a 2X enzyme and cofactor mixture in 1X reaction buffer and mixed with pipetting. 1 mM EDTA (final concentration) was used to quench the reactions at desired time points. The quenched reactions samples were then digested into single nucleosides with nuclease digestion (NEB nucleoside digestion mix, M0694S), using half of the recommended product volume and incubated overnight at 37 ^°^C. Samples with m^6^A_m_ capped substrates were first decapped with 1 µL of recombinant decapping enzyme (Edc1-Dcp1/Dcp2 complex stocked at 0.5 mg/ml) for 1 hour and then digested to single nucleosides as above.

### UHPLC-MS nucleoside analysis

After nucleoside digestion, the reaction samples with 1 µM FTO were precipitated by addition of 100% trichloroacetic acid to a final concentration of 20%. Samples were then centrifuged and supernatant was collected for LC-MS. Analysis of demethylation status was performed on an Agilent Bio-Inert 1260 Infinity II UHPLC system with Infinity Lab LC/MSD iQ. Separation of individual RNA nucleosides was conducted using an Agilent Zorbax SB-Aq Rapid Resolution HD (2.1 × 100 mm, 1.8 µm particle size) using mobile phase containing 0.1% formic acid (A) and 100% acetonitrile, 0.1% formic acid (B) and then detected by mass spectrometry in positive ionization mode. The gradient was as follows for detection of m6A at 0.480 mL/min (m/z = 282, 3.5 min) and A (m/z = 268, 1.6 min): 0-1.0 min, 100% A; 1.0 33 4.25 min, to 85% A/15% B; 4.25-5.0min, to 25% A/75% B; 5.0-7.0 min, 25% A/75% B; 7.0-7.1 min, to 100% B; 7.1-11.75 min, 100% B. The gradient was as follows for detection of m^6^A_m_ at 0.450 mL/min (m/z = 296, 3.65 min) and Am (m/z = 282, 3.5 min): 0-1.0 min, 100% A; 1.0-1.1 min, to 40% A/60% B; 1.1-5.0 min, to 28% A/72% B; 5.0-5.1 min, to 100% B; 5.1.0-8.0 min, 100% B.

### Protein digestion and LC-MS/MS analysis

Protein sample preparation was conducted as described previously with some modifications (Martin et al., 2024). 100 µL reactions were quenched with EDTA and then mixed with 4x volume of 80% acetonitrile. The quenched protein sample was then transferred to E3 filters (CDS analytics, Oxford, PA) and centrifuged at 400 x g for 2 mins. Filters were then washed with 200 µL of 80% acetonitrile followed by incubation at 45 ^°^C with 100 µL of 50 mM triethylammonium bicarbonate (TEAB), 10 mM Tris(2-carboxyethyl) phosphine (TCEP), 40 mM chloroacetamide (CAA) with gentle shaking. Filters were centrifuged and washed once with 200 µL of 80% acetonitrile. Protein digestion was performed with 200 µL of TEAB at trypsin:protein ratio 1:50 at 37 ^°^C for 18 hours with gentle shaking at 500 rpm.

Samples were then eluted with 0.1% formic acid in water followed by 50% acetonitrile/0.1% formic acid in water. The two elutions were pooled, vacuum dried, and desalted with C18 StageTips (CDS analytics, Oxford, PA). In brief, to activate the tips, 200 µL of 100% methanol was introduced and centrifuged at 1500 x g for 1 min followed by a second activation step of adding 80% acetonitrile/0.5% acetic acid. Tips were then equilibrated with 200 µL of 0.5% acetic acid. The dried peptides were resuspended in 200 µL equilibration buffer, briefly vortexed, and centrifuged at 16,000 x g for 5 mins before loading the tips. The tips were spun 400 x g for 10 mins and washed 3 times with 0.5% acetic acid in water. Peptides were eluted with 60% acetonitrile/0.5% acetic acid followed by 80% acetonitrile/0.5% acetic acid. Elutions were pooled, vacuum concentrated, and stored at -80°C until further analysis.

LC-MS/MS was performed on the Ultimate 3000 RSLC nano LC system coupled with Orbitrap Eclipse mass spectrometer and FAIMS Pro Interface (Thermo Scientific), and run as described previously (Martin et al., 2024). In brief, the dried peptides were first resuspended into 20 µL of LC buffer A (0.1% formic acid in water) and then loaded onto a trap column (PepMap100 C18,300 μm × 2 mm, 5 µm particle; Thermo Scientific) followed by separation on an analytical column (PepMap100 C18, 50 cm × 75 μm i.d., 3 μm; Thermo Scientific) at a flow of 250 nL/min. The LC gradient was configured as follows: 0-125 min, 1% to 25% LC buffer B (0.1% formic acid in ACN); 125-135 min, 25-32% LC buffer B; 135-140 min, 80% LC buffer B; 140-145, 80-1% B; 145-160 min, 1% B. The MS data were acquired under data-dependent acquisition (DDA) mode in Orbitrap at 60,000 resolution, followed by MS/MS acquisition of the most intense precursors for 1 s. For FAIMS, a 3-CV experiment (-40|-55|-75) was applied. The MS raw data were processed using MaxQuant and Andromeda software suite (version 2.5.2.0). Detailed settings include: Trypsin as digestion enzyme; methionine and leucine oxidation, acetyl (protein N-terminus) as variable modifications; cysteine carbamidomethylation as fixed modification; max missed cleavage 2; and minimum peptide length 7.

## ACKNOWLEDGMENTS

This work was supported by the US National Institutes of Health, National Institute of General Medical Sciences, under award R35 GM143000 to JSM, as well as P20 GM104316 that funded key instrumentation used in this study. The content is solely the responsibility of the authors and does not necessarily represent the official views of the National Institutes of Health. This work was further supported by grants 2019/33/B/ST4/01843 to JJ and 2022/47/D/ST4/00386 to MW from the National Science Centre, Poland.

## Notes

### Competing Interest Statement

The authors have declared no competing interest.

